# Photon-directed Multiplexed Enzymatic DNA Synthesis for Molecular Digital Data Storage

**DOI:** 10.1101/2020.02.19.956888

**Authors:** Howon Lee, Daniel J. Wiegand, Kettner Griswold, Sukanya Punthambaker, Honggu Chun, Richie E. Kohman, George M. Church

## Abstract

New storage technologies are needed to keep up with the global demands of data generation. DNA is an ideal storage medium due to its stability, information density and ease of readout with advanced sequencing techniques. However, progress in writing DNA is stifled by the continued reliance on chemical synthesis methods. The enzymatic synthesis of DNA is a promising alternative, but thus far has not been well demonstrated in a highly parallelized manner. Here, we report a novel multiplexed enzymatic DNA synthesis method using maskless photolithography. Rapid uncaging of Co^2+^ ions by patterned UV light activates Terminal deoxynucleotidyl Transferase (TdT) for spatially-selective synthesis on an array surface. Spontaneous quenching of reactions by the diffusion of excess caging molecules confines synthesis to light patterns and controls the extension length. We show that our multiplexed synthesis method can be used to store digital data by encoding 12 unique DNA oligonucleotide sequences with music from the 1985 Nintendo video game Super Mario Brothers™, which is equivalent to 84 trits or 110 bits of data.

---

In the era of Big Data, molecular DNA has become an increasingly attractive medium for the storage and archiving of digital data ^1–6^. This is primarily attributed to its ultra-high storage density, which is currently estimated to be in the hundreds of petabytes per gram DNA ^7^. The attractiveness of DNA as a data storage medium is additionally bolstered by its durability, longevity and energy efficiency compared to counterpart storage mediums; both analog and digital ^8,9^. However, for storing a meaningful volume of digital data, the synthesis of many unique DNA sequences at long lengths is required. While advances in array-based Oligonucleotide Library Synthesis (OLS) technology have enabled highly multiplexed DNA oligonucleotide synthesis, production in this format still relies on decades old phosphoramidite chemical synthesis methods ^10^. Many time-consuming steps of expensive and harsh reactions with the accumulation of toxic by-products greatly limit chemical synthesis as the demand for longer and larger quantities of DNA oligonucleotides increases ^11^.

Recently, the use of terminal deoxynucleotidyl transferase (TdT), a template-independent polymerase, to synthesize DNA oligonucleotides was shown to be a promising alternative to chemical synthesis ^12–14^. Because synthesis reactions are performed under aqueous conditions, many limiting aspects of the phosphoramidite chemistry can be circumvented or improved upon. This would be ideal for digital data storage in DNA; however, due to the natural promiscuity of TdT, controlling the enzyme in a sequence-specific manner is challenging ^15–17^. In order to overcome this challenge, a recent study showed controlled TdT extension activity with apyrase by degrading free nucleotides needed for synthesis ^14^. Incubation with optimized ratios of apyrase, TdT and nucleotide over multiple cycles resulted in the successful synthesis of several DNA oligonucleotides comprised of short homopolymeric blocks encoding digital information within unique base transitions. While there is a clear potential for enzymatic-based methods, its widespread adoption for applications such as digital data storage is hindered by the lack of parallelized synthesis technologies. Here, we demonstrate a novel enzymatic DNA oligonucleotide synthesis platform that utilizes photolithography to selectively modulate TdT extension activity in a highly multiplexed array format.

The photolithographic activation of TdT is achieved by optically controlling the ‘spatio-temporal’ concentration distribution of its metal cofactor cobalt (Co^2+^) near the array surface with the photocleavable caging molecule DMNP-EDTA ^18–20^. Initially, all Co^2+^ ions are complexed with DMNP-EDTA, which is provided in excess, keeping TdT in an inactive state until ready for synthesis. Upon photolysis of complexed DMNP-EDTA with patterned UV light, Co^2+^ ions are uncaged resulting in a localized pulse of cofactor concentration and spatially selective oligonucleotide extension by TdT within the defined areas of irradiation. To control the number of extension events, the UV light source is turned off after a predetermined time allowing excess DMNP-EDTA to diffuse into irradiated areas, chelating Co^2+^ ions and returning TdT to an inactive state. This sequence of events constitutes a full cycle of synthesis **(Fig. 1a)**. Virtually any individual position on the array surface is spatially addressable by our system using a computer-controlled spatial light modulator, or digital micromirror device (DMD), to facilitate the on-demand generation of dynamic light patterns without needing to construct physical masks **(Fig. 1b)**. This functionality is paramount for controlled multiplexed oligonucleotide synthesis on the array surface.

**Figure 1.**
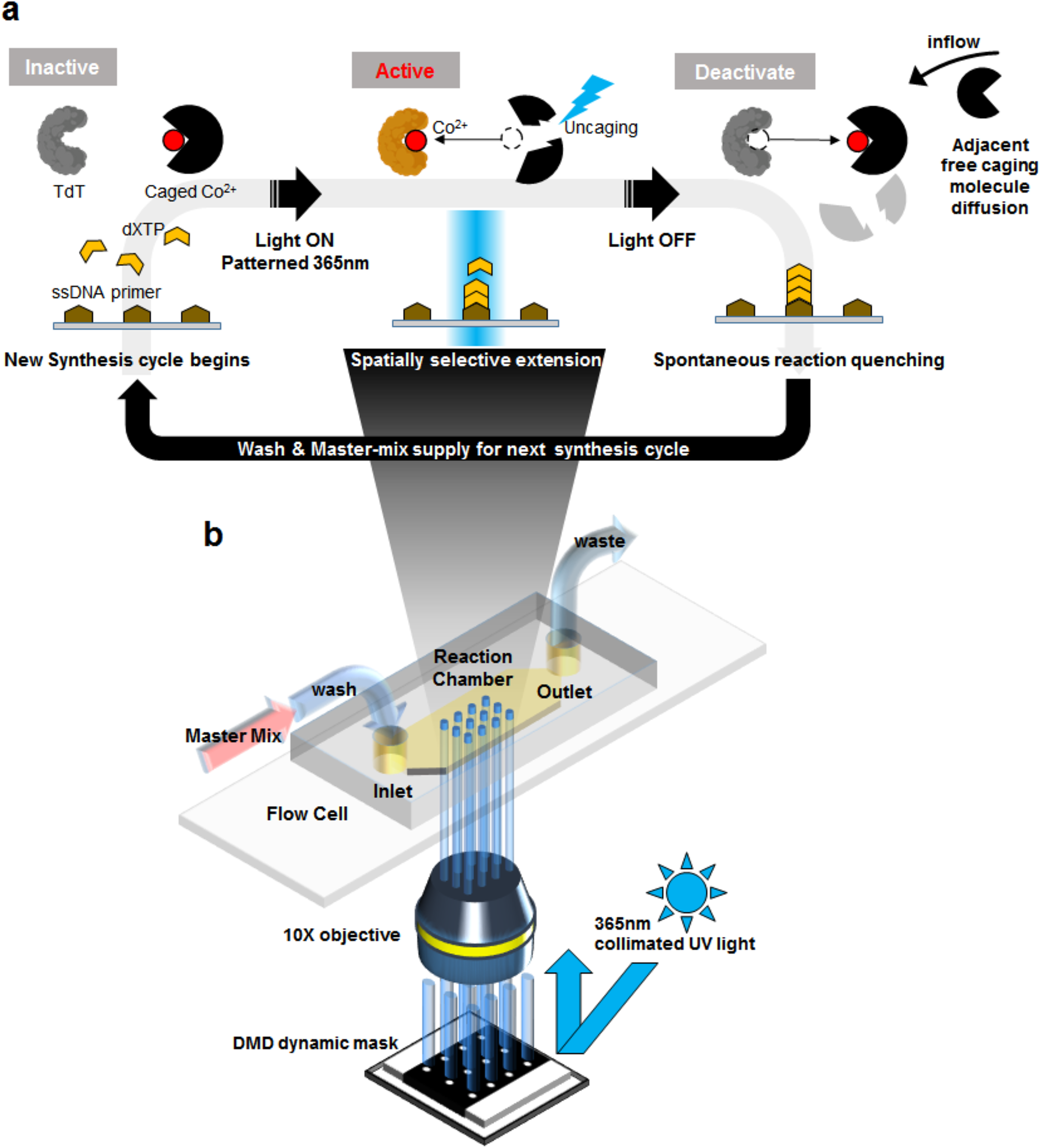
Overview of Photon-directed Multiplexed Enzymatic DNA Synthesis System. **a**, An array surface derivatized with single-stranded DNA initiator oligonucleotide is brought into contact with a master mix containing the appropriate buffers, Co^2+^ divalent cation cofactor, TdT enzyme, the desired nucleotide to be incorporated (dXTP) and photolabile DMNP-EDTA caging molecule provided in excess. All Co^2+^ ions are initially complexed with DMNP-EDTA, which causes TdT to remain in an inactive state until needed *(Inactive)*. Using photolithography, patterned UV light at 365 nm illuminates the array’s surface causing the complexed DMNP-EDTA to degrade releasing Co^2+^ ions and activates TdT in a spatially selective manner *(Active)*. The UV light is then turned off and the reaction is allowed to incubate for a short period of time. During this incubation, excess, non-complexed DMNP-EDTA spontaneously quenches the extension reaction by chelating free Co^2+^ causing active TdT to become inactive *(Deactivate)*. The array surface is then washed and either the next synthesis cycle begins or material is retrieved from the surface for downstream applications. **b,** Arrays are mounted into a simple flow cell with a reaction chamber inlet and outlet to waste or collection. Individually addressable patterning is a major advantage of our synthesis method, which is provided by the generation of reflective on-demand dynamic masks from the DMD through a 10X objective from a collimated UV light source.

We initially found that several key system parameters required optimization before we could perform template-independent DNA oligonucleotide synthesis in multiplex. First, irradiation with UV light at an energy density of at least 2J/cm^2^ (2W/cm^2^ for 8-15 seconds) was critical for the sufficient uncaging of Co^2+^ ions and TdT activation. Lower UV doses resulted in little observable enzymatic activity (data not shown), suggesting that either incomplete photolysis of DMNP-EDTA occurred or that the uncaging rate of Co^2+^ ions was significantly lower than the rate at which they were spontaneously chelated by free DMNP-EDTA. Supporting evidence for the latter was given by the necessity to finely tune the total concentration of DMNP-EDTA initially supplemented in the synthesis reaction master mix.

We determined that the DMNP-EDTA concentration needed to be at least 1.3X that of Co^2+^ for proper spatial confinement on the surface; however, this was also largely dependent on the fill factor of the pattern generated by the DMD (**Supplementary Fig. 1**). We also found that a post-illumination incubation for 15 to 20 seconds significantly boosted the overall yield of oligonucleotide extension on the surface **(Supplementary Fig. 2)**. This incubation period allowed TdT enough time to incorporate nucleotides upon Co^2+^ ion uncaging before being quenched by DMNP-EDTA when the UV light source was turned off. The enzyme concentration and percentage of glycerol in the master mix also affected the kinetics of nucleotide incorporation, but were overall less impactful **(Supplementary Fig. 3)**. All reactions were performed at room temperature rather than 37 °C for user convenience and to prevent excessive extension by TdT.

While optimization of our system was multifactorial, we ultimately decided to use a mask pattern of 100 μm diameter circular spots arranged in a (3 × 4) array on 1.2 mm^2^ of surface area. This yielded 12 individually addressable locations for multiplexed oligonucleotide synthesis **(Fig. 2a)**. To confirm that these conditions were suitable for proper extension, we synthesized homopolymers consisting of only “G” at all 12 spots from the 3’-terminus of short initiator sequences anchored to the surface. Positive extension was verified using fluorescent imaging after the ligation of a short visualization probe containing a Cy3 fluorophore using a sequence specific splint (**Supplementary Fig. 4a, Methods).** No extension outside the illuminated areas or erroneous crosstalk between the 12 spots was observed on the surface, indicating robust nucleotide incorporation by TdT and that DMNP-EDTA quenching functioned appropriately to spatially confine DNA oligonucleotide synthesis **(Fig. 2b)**.

**Figure 2.**
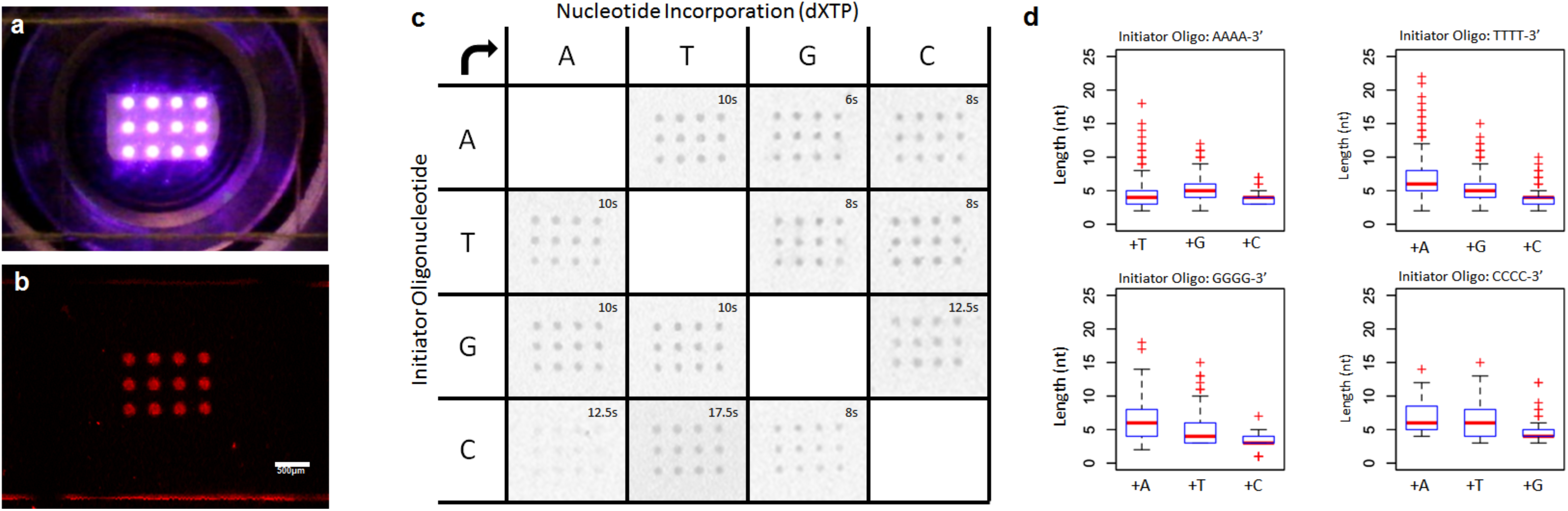
Multiplex Enzymatic Synthesis Optimization and Base Transition Normalization. **a**, Demonstration of UV irradiation of the array surface with 100 μm circular spots arranged in a (3 × 4) patterned format on 1.2 mm^2^ of surface area. UV irradiation is not limited to this particular patterning and may be pixel-wise (1920×1080) changed on-demand with our photolithographic system. Any spot on the surface is individually addressable in terms of spatial location and the total amount of UV irradiation time. **b,** Visualization of “G” homopolymeric oligonucleotide synthesis post system optimization via the splint-end ligation of a probe sequence containing a 3’-Cy3 fluorophore using the (3 × 4) pattern. **c,** Results of base transition normalization in which the total illumination time was adjusted for any base transition that may be encountered during multiplexed synthesis. The left axis indicates the composition of the last 4 bases on the 3’-terminus of surface initiator oligonucleotide and the top axis indicates the nucleotide that was incorporated onto the respective initiator oligonucleotide. The optimal illumination time is indicated in each base transition box and was determined by splint-end ligated 3’- Cy3 fluorescent signal. **d,** Box plots indicating NGS analysis of base transition normalization. Graphs show the normalized nucleotide (nt) extension length distribution for all possible base transitions with red pluses being statistical outliers.

We next sought to normalize the rate of incorporation for all nucleotide types by TdT in preparation for multi-cycle oligonucleotide synthesis as it is well-known that they can be highly variable given the composition of the initiator sequences’ last 3-5 bases ^16^. To do this, we increased or decreased the total UV illumination time on the surface for a given base transition. Using the same 12 spot pattern, we empirically found that base transitions involving initiator oligonucleotides ending with “G” or “C” required longer illumination times (>12 seconds) as compared to those ending with “A” or “T” (≤10 seconds) for all nucleotide types **(Fig. 2c)**. The longest illumination time was 17.5 seconds for the base transition “C” to “T”, while the shortest was 6 seconds for “A” to “G”.

Because these results only indicated a relative comparison of overall TdT activity for each condition, we employed next-generation sequencing (NGS) to determine the actual number of extension events that occurred for each base transition. We again utilized our sequence specific splint-end ligation approach to ligate the appropriate adaptors for PCR amplification and NGS library preparation. With this, only those sequences that were properly and robustly extended would be extracted for NGS analysis (**Supplementary Fig. 4b)**. Overall, we found that the normalization of illumination times resulted in an average of 4 to 8 extension events for all tested base transitions; however, >15 extension events were possible, but were only observed in a fraction of what was synthesized (**Fig. 2d)**.

We next sought to prove that our system was capable of multi-cycle synthesis by producing a heteropolymeric sequence consisting of all four natural nucleotide bases in single-plex on the (3 × 4) array **(Fig. 3a)**. We elected to synthesize the 8 base transition DNA sequence “GATGTAGAC” with the expectation that a successful demonstration would yield oligonucleotide comprised of short homopolymeric blocks for each base used, similar to previous work ^14^. Synthesis began with an anchored initiator oligonucleotide consisting of a 3’- string of four “G’s” and ended with a final base transition of “C” for adaptor ligation and sequencing purposes. Enzyme master mixes supplemented with the appropriate nucleotide were applied to the flow cell in the designated order with all 12 spots illuminated during each cycle to produce the target sequence. Following the completion of 8 synthesis cycles, we verified that the correct sequence was produced and determined the extension length distribution for each homopolymer block using NGS as previously outlined **(Fig. 3b**, **Methods)**. We additionally tracked the progress after each cycle by performing parallelized synthesis across several flow cells with which we could then analyze TdT extension with gel electrophoresis after retrieval from the surface **(Fig. 3c)**. In both cases, the distribution of block lengths were in line with what we expected post-normalization of the illumination times.

**Figure 3.**
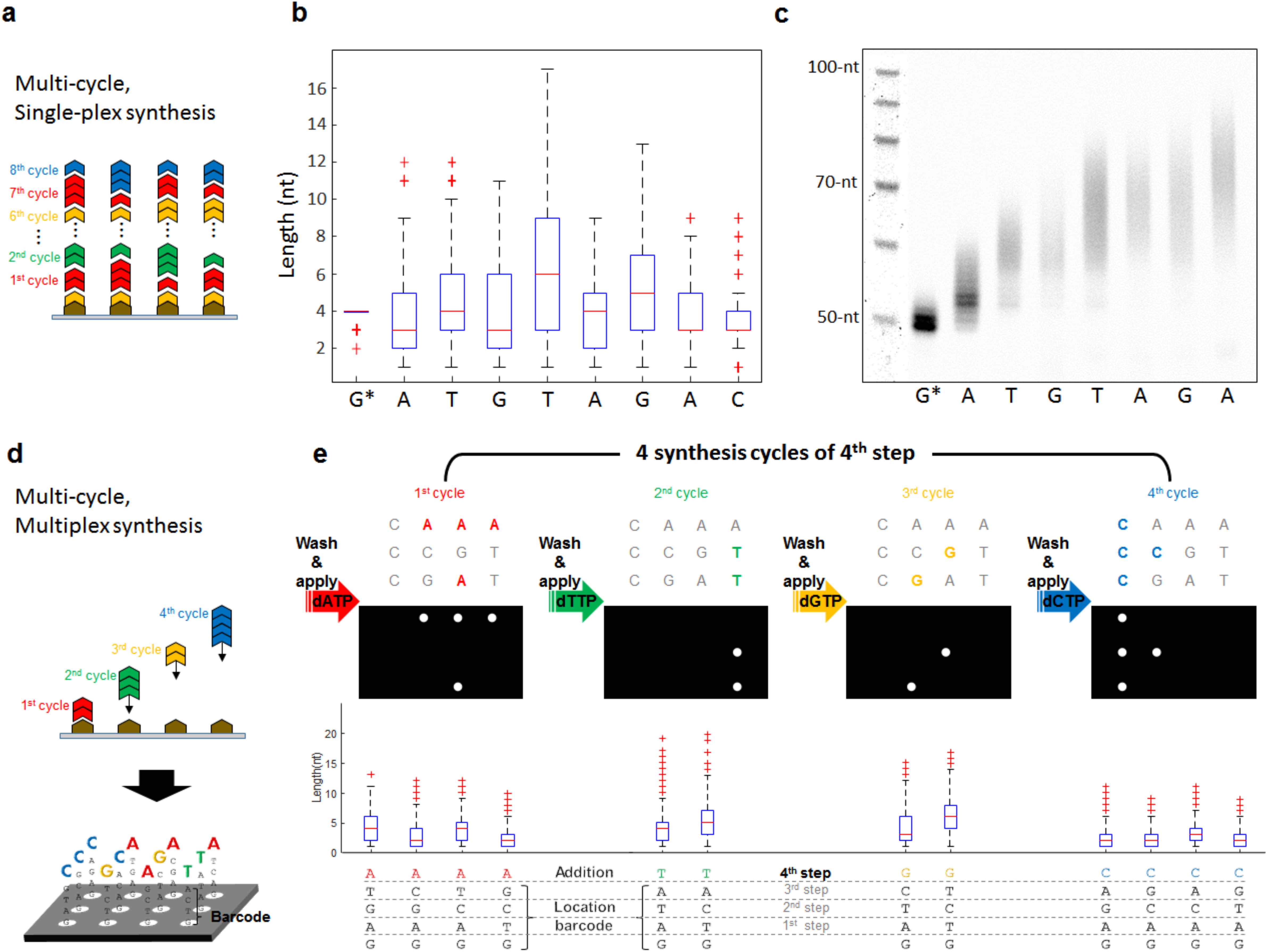
Demonstration of Multi-cycle and Multiplex Enzymatic Synthesis. **a**, An overview of multi-cycle, single-plex synthesis of a heteropolymer oligonucleotide comprised of 8 unique base transitions. Each cycle represents the addition of a single nucleotide base type at all 12 spots on the (3 × 4) array. Synthesis results in short homopolymeric blocks of A, T, G or C at variable lengths. **b,** Following multi-cycle synthesis and NGS, raw sequence reads were extracted and filtered by the presence of the adaptor sequence added by splint-end ligation after the final “C” extension. The box-plot indicates a statistical representation of the number of extension events for each homopolymeric block for the synthesized sequence “GATGTAGAC”. Note that only sequencing reads that contained all 8 base transitions were used to generate the box-plot and that synthesis started with a string of four “G” on the 3’-terminus of the initiator oligonucleotide. **c,** Denaturing gel electrophoresis analysis of each individual cycle of synthesis. Each lane represents the material retrieved of an individual flow cell after the appropriate number of cycles were performed. No final “C” extension or splint-end ligation occurred for this analysis. **d,** An overview of multi-cycle, multiplex synthesis of 12 unique heteropolymer oligonucleotides on the (3 × 4) array. Each synthesis step contained the individual cycles for the addition A, T, G, and C at the appropriate spots on the array. Sequence barcodes indicating the physical location of each oligonucleotide on the array can be built into the initiator oligonucleotide or synthesized by TdT. For each cycle, a unique mask was generated by the system’s DMD to locally activate TdT based on the desired sequence to be synthesized. **e,** For example, the masks needed for the 4th step of an 8-step multiplex oligonucleotide synthesis run is shown with the appropriate post-synthesis sequencing data to verify spatially-selective synthesis.

Having determined that multi-cycle synthesis was possible in single-plex, we next turned to showing that the same process could be applied in multiplex in order to produce 12 unique oligonucleotide sequences on the array surface simultaneously **(Fig. 3d)**. We accomplished this by taking advantage of the ability for our system’s DMD to rapidly generate dynamic mask patterns on demand. This allowed us to incorporate nucleotide bases at any particular spot or combination of spots on the (3 × 4) array. As an added benefit, individual spot illumination time could be adjusted to meet the established TdT normalization criterion for any given base transition **(Fig. 2c, d)**. A total of 29 unique masks and incorporation cycles over 8 steps of synthesis were required to produce the desired oligonucleotides in multiplex **(Supplementary Fig. 5)**. One step of multiplex synthesis contained the incorporation cycles for all four natural nucleotides, which could be algorithmically filtered and individually analyzed for homopolymeric block length post-sequencing **(Fig. 3e)**. If a step did not require one or more bases, those cycles were skipped by not illuminating the surface.

To demonstrate that our synthesis method can be used for digital data storage applications, the 12 oligonucleotide sequences synthesized in multiplex encoded the first two measures of the “Overworld Theme” sheet music from the 1985 Nintendo Entertainment System (NES) video game Super Mario Brothers™ **(Supplementary Fig. 6a, b).** Our encoding and decoding scheme was based on previous work ^14^, where digital data is stored into the homopolymeric blocks of unique base transitions produced by TdT **(Fig. 4a-g)**. Input musical information was indexed, assigned a note number based on a Musical Instrument Digital Information (MIDI) chart, converted to ternary, and from this, the 12 unique DNA oligonucleotide sequences were generated **(Fig. 4a-c**, **Supplementary Fig. 7a-c, 8)**. In addition to sheet music, which contained information regarding the notes as well as the order and duration each note is to be played, we used the first three base transitions to store the physical locations of each oligonucleotide on the (3 × 4) array as barcodes. In total, this amounted to 84 trits, or 110 bits, of data stored digitally in the 12 DNA oligonucleotide sequences.

**Figure 4.**
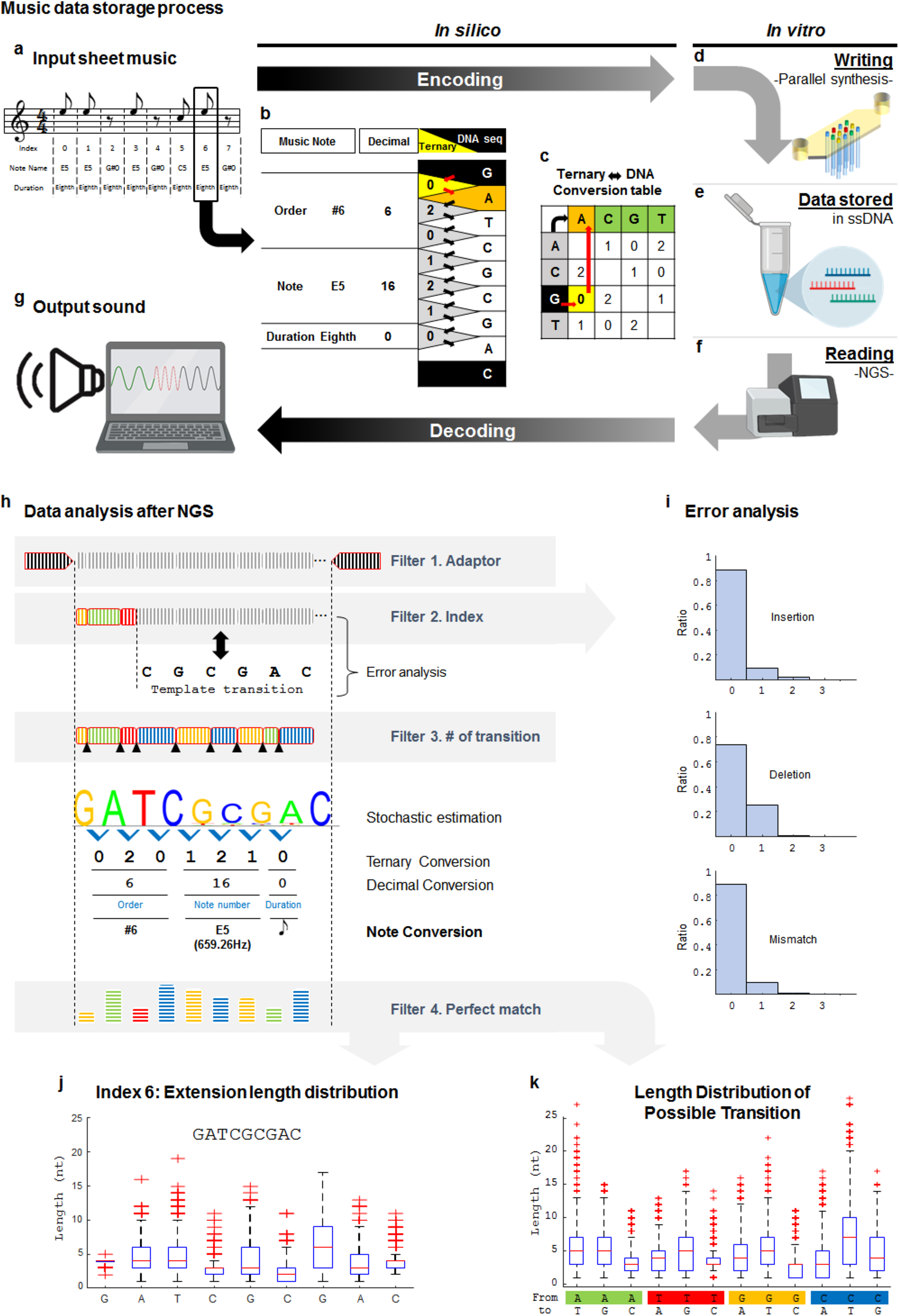
Demonstration of Digital Music Data Storage in DNA Oligonucleotides Enzymatically Synthesized in Multiplex. **a**, A snapshot of the simplified melody from the two first measures of the 1985 Nintendo Entertainment System video game Super Mario Brothers™ “Overworld Theme” piano sheet music. Each note and rest of the melody was indexed as #0 through #11 and stored into one of the 12 unique DNA oligonucleotides synthesized in multiplex on the (3 × 4) array. The full piano sheet music is shown in **Supplementary Fig. 6**. **b,** Overview of the *in silico* encoding and decoding scheme for the storage of digital music into DNA. Indexed notes and rests were assigned a note number based on a modified Musical Instrument Digital Information note chart, which also indicates note octave. In addition to the note, the encoding scheme also stores the order and duration each note is to be played. Note and rest indices double as the physical locations of their respective oligonucleotides on the (3 × 4) array and the play order. All rests are assigned to the note G#_0_, which is inaudible (25.9 Hz) to normal human hearing ^22^. A full overview of MIDI conversion of sheet music to digital data is shown in **Supplementary Fig. 7**. **c,** Digital music data is translated to ternary and the unique DNA sequences are mapped to unique base transitions using a conversion map. The left axis of the table represents the string of bases at the 3’- terminus of the initiator oligonucleotide and the top axis indicates the nucleotide to be incorporated next. **Supplementary Fig. 8** indicates a table outlining the mapped “Overworld Theme” musical melody to DNA sequences with all relevant data and information. **d,** Generated DNA sequences are enzymatically synthesized with TdT using our multiplex photolithographic system *(Writing)*. **e,** Synthesized oligonucleotides are retrieved from the flow cell surface and stored in tubes for sequencing or other downstream applications *(Storage)*. **f,** To read and decode stored digital music data, DNA oligonucleotides synthesized in multiplex can be sequenced with NGS such as Illumina or nanopore methodologies *(Reading)*. **g,** From sequencing data, the decoding process converts sequencing information back to musical notes. A sinusoidal wave generator is used to play the “Overworld Theme” melody in the correct note order with the proper duration in true sound. **h,** During the decoding process, several filters are applied to extract and align the reads that contain the digital music data to the expected template sequences. Template alignment was performed using the Smith-Waterman algorithm. **i,** From this, error analysis can be performed to determine the quality of multiplex synthesis. Bar graphs indicate the percentage of insertions, deletions or mismatches that occurred in the filtered sequencing reads. **j.** Sequencing data also yields statistical information regarding the extension length distribution for each base transition for all 12 oligonucleotides synthesized in multiplex. For example, subset 6, which indicates sequence index 6 is shown in the box plot. All other subsets are indicated in **Supplementary Fig. 9, 10. k**, Additional statistics such as the extension length distribution for all possible transitions from the entire array were analyzed as shown in the indicated box plot.

Following multiplex synthesis and NGS, filtered sequence data was algorithmically decoded and processed by a sinusoidal wave generator to play the sheet music from in true sound **(Supplementary File 1)**. Filtering occurred in four discrete stages **(Fig. 4h)**. We first selected reads that contained the correct initiator oligonucleotide and splint-ligated sequencing adapter. From these, we subsetted those with 1 of the 12 index barcodes and then selected for sequences that indicated 8 full base transitions within the individual subsets. Stochastic estimation indicated that the dominant population of each subset was perfectly matched to their respective template sequences without needing to implement additional error correction bits **(Fig. 4j**, **Supplementary Fig. 9a-c, 10a,c)**. We found that each subset contained the expected normalized distribution of homopolymeric block extension lengths. This was also observed across the entire (3 × 4) array for all possible base transitions during multiplex synthesis **(Fig. 4k**, **Supplementary Fig. 10b)**. A nominal portion of the filtered sequences contained errors despite our optimization efforts. These errors most likely arose from incomplete washing of the flow cell, unwanted crosstalk between spots (insertion 11.6% or mismatch 10.1%) or insufficient activation of TdT (deletion 26.1%) **(Fig. 4i)**.

Taken together, our results represent the first demonstration of template-independent, multiplexed enzymatic synthesis of DNA oligonucleotides for digital data storage through the application of photolithography to control enzymatic polymerization. Our system utilizes a combination of commercial photolabile materials and maskless lithography technology with the added benefit of circumventing the use of expensive physical photomasks, special phosphoramidites, harsh organic solvents and the accumulation of toxic waste from chemical synthesis. Additionally, since metal cation cofactors are essential for polymerase catalysis, the presented method will be potentially suitable for controlled synthesis using mutated enzymes with enhanced attributes such as higher processivity or the ability to incorporate modified nucleotides ^21^. While enzyme-based oligonucleotide synthesis technology is still in its infancy, we envision that bridging overall cleaner and faster DNA oligonucleotide synthesis with cutting-edge, high-density array photolithographic techniques will pave the way for its widespread adoption and further promote practical molecular data storage technology.

## Code Availability

Encoding and decoding MatLab scripts used in this study are available at: http://github.com/dwiegand740/Photon_Enzymatic_Synthesis

## Materials & Methods

See supplementary information for all materials and methods.

## Acknowledgements

The authors would like to thank Dr. Henry Lee for useful discussions regarding digital data storage, enzyme kinetics and next-generation sequencing. The authors like to additionally thank Mr. Benjamin Vieira for the helpful discussion of musical nomenclature. K.G. is supported under Graduate Fellowships from the Fannie and John Hertz Foundation and the Charles Stark Draper Laboratory.

## Author contributions

HL, KG, & GMC conceptualized the project. HL, DJW, KG, SP and HC conducted the experiments involving: design and initiation of DNA synthesis (HL, DJW and KG), digital music encoding (HL), maskless photolithography system and flow cell development (HL and HC) and sequencing & analysis (HL, DJW and SP). HL, DJW, SP, REK, & GMC wrote the manuscript. REK & GMC supervised the study.

## References

1. Church, G. M., Gao, Y. & Kosuri, S. Next-generation digital information storage in DNA. Science 337, 1628 (2012).

2. Goldman, N. et al. Towards practical, high-capacity, low-maintenance information storage in synthesized DNA. Nature 494, 77–80 (2013).

3. Ceze, L., Nivala, J. & Strauss, K. Molecular digital data storage using DNA. Nat. Rev. Genet. 20, 456–466 (2019).

4. Koch, J. et al. A DNA-of-things storage architecture to create materials with embedded memory. Nature Biotechnology 1–5 (2019).

5. Anavy, L., Vaknin, I., Atar, O., Amit, R. & Yakhini, Z. Data storage in DNA with fewer synthesis cycles using composite DNA letters. Nat. Biotechnol. (2019) doi:10.1038/s41587-019-0240-x.

6. Organick, L. et al. Probing the physical limits of reliable DNA data retrieval. Nat. Commun. 11, 616 (2020).

7. Erlich, Y. & Zielinski, D. DNA Fountain enables a robust and efficient storage architecture. Science 355, 950–954 (2017).

8. Bancroft, C., Bowler, T., Bloom, B. & Clelland, C. T. Long-term storage of information in DNA. Science 293, 1763–1765 (2001).

9. Zhirnov, V., Zadegan, R. M., Sandhu, G. S., Church, G. M. & Hughes, W. L. Nucleic acid memory. Nat. Mater. 15, 366–370 (2016).

10. Kosuri, S. & Church, G. M. Large-scale de novo DNA synthesis: technologies and applications. Nat. Methods 11, 499–507 (2014).

11. LeProust, E. M. et al. Synthesis of high-quality libraries of long (150mer) oligonucleotides by a novel depurination controlled process. Nucleic Acids Res. 38, 2522–2540 (2010).

12. Palluk, S. et al. De novo DNA synthesis using polymerase-nucleotide conjugates. Nat. Biotechnol. 36, 645–650 (2018).

13. Barthel, S., Palluk, S., Hillson, N. J., Keasling, J. D. & Arlow, D. H. Enhancing Terminal Deoxynucleotidyl Transferase Activity on Substrates with 3’ Terminal Structures for Enzymatic De Novo DNA Synthesis. Genes 11, (2020).

14. Lee, H. H., Kalhor, R., Goela, N., Bolot, J. & Church, G. M. Terminator-free template-independent enzymatic DNA synthesis for digital information storage. Nat. Commun. 10, 2383 (2019).

15. Moon, A. F. et al. The X family portrait: structural insights into biological functions of X family polymerases. DNA Repair 6, 1709–1725 (2007).

16. Motea, E. A. & Berdis, A. J. Terminal deoxynucleotidyl transferase: the story of a misguided DNA polymerase. Biochim. Biophys. Acta 1804, 1151–1166 (2010).

17. Chang, L. M. S., Bollum, F. J. & Gallo, R. C. Molecular Biology of Terminal Transferas. Crit. Rev. Biochem. Mol. Biol. 21, 27–52 (1986).

18. Adams, S. R. & Tsien, R. Y. Controlling cell chemistry with caged compounds. Annu. Rev. Physiol. 55, 755–784 (1993).

19. Ellis-Davies, G. C. R. Caged compounds: photorelease technology for control of cellular chemistry and physiology. Nat. Methods 4, 619–628 (2007).

20. Ji, G., Feldman, M., Doran, R., Zipfel, W. & Kotlikoff, M. I. Ca2+ -induced Ca2+ release through localized Ca2+ uncaging in smooth muscle. J. Gen. Physiol. 127, 225–235 (2006).

21. Jensen, M. A. & Davis, R. W. Template-Independent Enzymatic Oligonucleotide Synthesis (TiEOS): Its History, Prospects, and Challenges. Biochemistry 57, 1821–1832 (2018).

22. Rosen, S. & Howell, P. Signals and Systems for Speech and Hearing. (BRILL, 2011).

