## Supplementary Information for "Photon-directed Multiplexed Enzymatic DNA Synthesis for Molecular Digital Data Storage"

### Materials & Methods

#### Maskless Lithography System & Flow Cell Fabrication

Our maskless lithography system consisted of a high-power UV LED array with collimation adaptor (LumiBright PR, 2910A-100, Innovations in Optics) couple with Tube lens (MT-L Accessory Tube Lens, Edmund Optics), DMD (DLP4000, Texas Instruments), and Objective lens (CFI Plan Fluor 10X, Nikon) to expose an ultraviolet (365 nm) pattern **(Fig. 1a, b)**. A self-designed computer program in MatLab™ and control circuit board (Arduino Uno) was used to synchronize DMD patterning and UV illumination time. Flow cells consisted of a cover, spacer and bottom glass slide with a single inlet and outlet **(Fig. 1b)**. The cover and spacer were fabricated out of acrylic and assembled with double-sided adhesive tape (9172MP, 3M). The inlet and outlet on the flow cell cover and the flow cell pattern of the spacer was precisely cut using a laser cutter (Epilog Legend 36EXT).

#### Initiator Oligonucleotide Surface Derivatization & Sequence Composition

A streptavidin-coated glass slide (ArrayIt Corporation) was used to anchor 5'- biotinylated initiator oligonucleotide (Integrated DNA Technologies), which served as the bottom of the fully assembled flow cell. To derivatize the streptavidin-coated glass slide, biotinylated oligonucleotide was incubated at a final concentration of 0.25 mM in 1X Binding & Wash buffer (20 mM, 1M NaCl, 1 mM EDTA, 0.0005% Triton-X100, pH 7.5) for 1 hour at room temperature. After incubation, the surface was washed with fresh 1X Binding & Wash buffer and then washed again with 1X phosphine buffer saline (PBS). A standard initiator oligonucleotide consisted of the sequence (/5Biosg/TGGTTAGTGTGCTTCGGACCGGGG) for initial system optimization and final multiplexed synthesis demonstration. For normalized base transition experiments, the initiator oligonucleotide sequence was the same with the exception of the last four bases on the 3'-end of the oligonucleotide. These were variable (either -GGGG, -CCCC, -AAAA, or -TTTT) depending on the target base transition.

#### Standard Master Mix Composition

A standard synthesis master mix was composed of 20 units of recombinant calf thymus TdT enzyme (Thermo Scientific), 1X reaction buffer (0.2 M potassium cacodylate, 0.025 M Tris, 0.01% (v/v) Triton X-100, 1 mM $CoCl_2$), 0.1 mM of either dATP, dTTP, dGTP, or dCTP (Invitrogen), and 1.3 mM caging molecule DMNP-EDTA tetrapotassium salt (MilliporeSigma) in 10 µL of deionized water. All reaction incubations were performed at room temperature.

#### Standard Synthesis Cycle

A standard synthesis cycle consisted of loading the necessary mask image to the DMD, delivering 2 µL of synthesis master mix to the flow cell, and then applying UV irradiation to the bottom surface for 10-20 seconds depending on the base transition occurring. A post-illumination incubation would then take place for a minimum of 20 seconds for optimal TdT extension and reaction quenching. Synthesis master mix would then be removed from the flow cell to the waste by vacuum and then washed with 20 µL of 1X PBS. A new mask image would be loaded into the DMD and this process would be repeated for all nucleotides to be incorporated during the synthesis cycle.

**Sequence-specific Split-end Ligation**

To visualize oligonucleotide synthesis without removing oligonucleotide from the surface, we employed a sequence-specific splint-end ligation technique [1] that allowed us to add a short oligonucleotide probe containing a 3'- Cy3 fluorophore to the end any oligonucleotide synthesized using our method **(Supplementary Fig. 4a, b)**. Splint-end ligations were performed using a Quick Ligation Kit (NEB) as per manufacturer's instructions. Reactions consisted of 1x Quick Ligase Buffer, 25 µM Cy3 labeled probe (/5Phos/CGA CTG AAC CCA AGC AAC TGA/3Cy3Sp/), 20 µM splint oligonucleotide (CA GTT GCT TGG GTT CAG TCG XXXX, X can be A, G, T, C depending on the synthesized strand), and 1 µL of Quick Ligase in deionized water in 20 µL of volume. 5 µL of the Quick Ligation master mix was delivered to the flow cell and incubated for 2 hours at 16 °C. After incubation, the flow cell was washed, disassembled, dried thoroughly with forced air, and imaged using a Typhoon FLA 9000 Imager (GE) with settings for Cy3 fluorescence filter with a 10 µm resolution. This ligation technique was additionally applied to selectively add PCR amplification and NGS adaptors to oligonucleotides correctly synthesized.

**Retrieval of Oligonucleotides from Surface**

To retrieve surface-bound oligonucleotide post-synthesis, a solution of 95% formamide in deionized water supplemented with 10 mM EDTA was applied to the disassembled flow cell bottom surface and heated to 65 °C for 5 minutes. This solution was then removed from the flow cell bottom surface and cleaved oligonucleotides were purified using a Clean & Concentrator Oligonucleotide Spin Column (Zymo) as per manufacturer's instructions. Oligonucleotide were eluted into deionized water for downstream processing or sequencing.

**PCR Amplification & Gel Electrophoresis Analysis**

Oligonucleotides removed from the surface with appropriate adaptors for PCR were amplified using a KAPA SYBR Fast qPCR 2X Master Mix Kit (Roche) as per manufacturer's instructions and purified QiaQuick PCR Clean-up columns (Qiagen). Amplified or raw oligonucleotide material was analyzed using 15% TBE-Urea precast polyacrylamide gels as per manufacturer's instructions. Oligonucleotide material amplified with Cy3 labeled primers was visualized using a Typhoon FLA 9000 Imager (GE) with settings for Cy3 fluorescence following gel electrophoresis.

**Illumina MiSeq Library Preparation & Sequencing**

Purified PCR amplified sequences containing encoded data were prepared for next-generation sequencing using a NEBNext Ultra II DNA Library Prep Kit for Illumina as per manufacturer's instructions with approximately 100 ng of material per library. Libraries were not size-selected during magnetic bead clean-ups in order to preserve the true length distribution of sequences synthesized on the surface of the arrays as best as possible. Libraries were indexed using a NEBNext Singleplex primer set. The extent of sequencing ligation and library indexing was then quantified using a NEBNext Library Quant Kit for Illumina as per manufacturer's instructions. Quantified libraries were then combined at an equimolar ratio for next-generation sequencing using an Illumina MiSeq sequencer.

**Oxford Nanopore Library Preparation & Sequencing**

For nanopore sequencing via the Oxford Nanopore technology method, purified PCR amplified sequences containing encoded data were subjected to library preparation using the 1D Genomic DNA ligation sequencing kit (SQK-LSK109) from Oxford Nanopore Technologies following the manufacturer's protocols. Briefly, 0.5 pmol of the double stranded DNA strands were used as starting material. The DNA was repaired and end-prepped using NEBNext FFPE DNA repair mix (NEB, M6630S) and NEBNext Ultra II End Repair/dA-Tailing Module (NEB, E7546) followed by bead purification using Agencourt AMPure XP beads (Beckman Coulter, A63880) at 1:2 sample to bead ratio. Adapters were then ligated to the end-prepped samples using the NEBNext Quick T4 DNA ligase (NEB, E6056S). The flow cells (R4.2.1) were primed, the sample was loaded onto the priming port of the flow cell and sequenced on the MinION, that generated about 500K reads/hr. Sequencing was performed using the MinKNOW software (version 18.3.1, Oxford Nanopore Technologies) that converted raw data (in the form of fast5 files) into fastQ files which were used for downstream analysis.

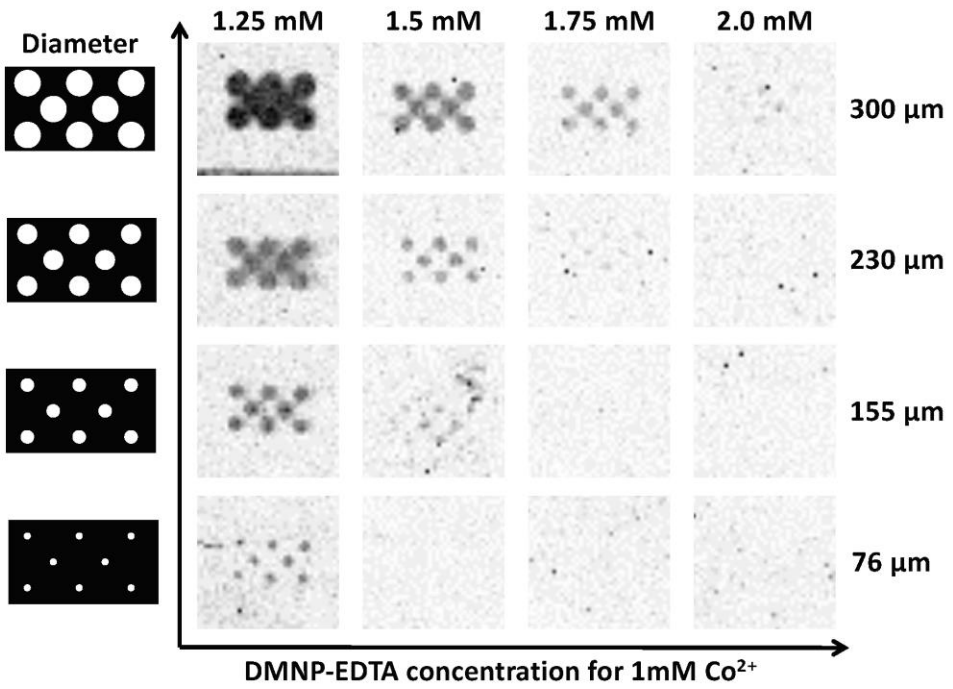


**Supplementary Figure 1:** A comparison of DMD pattern fill factor and concentration of the caging molecule, DMNP-EDTA. The fill factor is defined as the ratio between the illuminated and surrounding area within the pattern. For circular spots, the fill factor is dictated by their diameter. A finely tuned balance between a pattern's fill factor and caging molecule concentration present in the reaction master mix is required for well-confined nucleotide extension. This is indicated by sharp patterning and no cross-talk between the individual spots post-synthesis. As the fill factor (diameter of the spots) increases, less DMNP-EDTA is present in the surrounding area to chelate free $Co^{2+}$ that diffuses away from the illuminated pattern. Increasing the total concentration of DMNP-EDTA will help eliminate significant cross-talk between spots; however, too much DMNP-EDTA will quench synthesis reactions before visualible oligonucleotide extension can occur. Balanced fill factor and DMNP-EDTA is dependent on the total concentration of $Co^{2+}$ initially supplemented in the synthesis master mix. Optimization took place with 1 mM initial $Co^{2+}$, 8 seconds of UV irradiation, and 15 seconds of post-illumination incubation for all conditions.

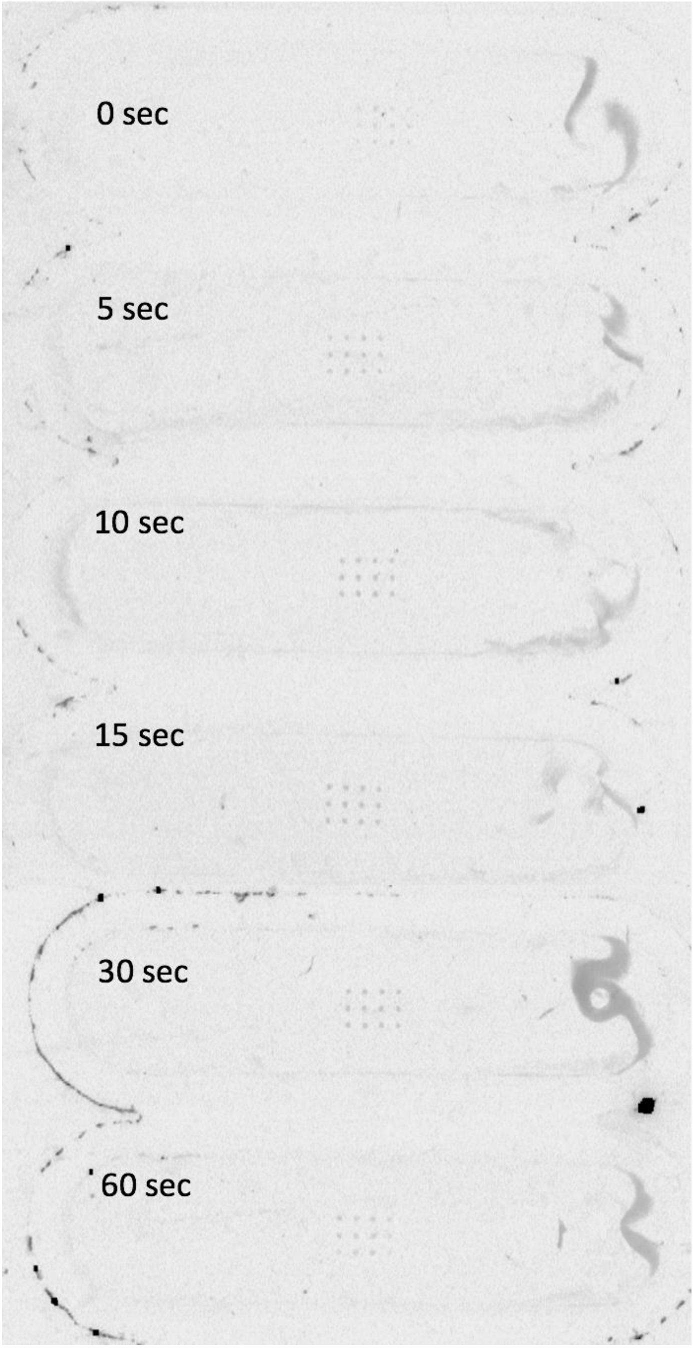


**Supplementary Figure 2:** A comparison post-illumination incubation times. It is essential to allow sufficient time for TdT to incorporate nucleotides onto the surface bound oligonucleotide before washing away the enzyme master mix containing free $Co^{2+}$ released from photolabile DMNP-EDTA upon UV irradiation. Using sequence-specific splint-end ligation with short visualization pro containing a 3'-Cy3 fluorophore, it was found that at least 5 seconds of post-illumination incubation is required for high-quality surface extension. Since DMNP-EDTA is supplemented in the enzyme master mix in excess, there were no significant differences between post-illumination incubation times longer than 15 seconds.

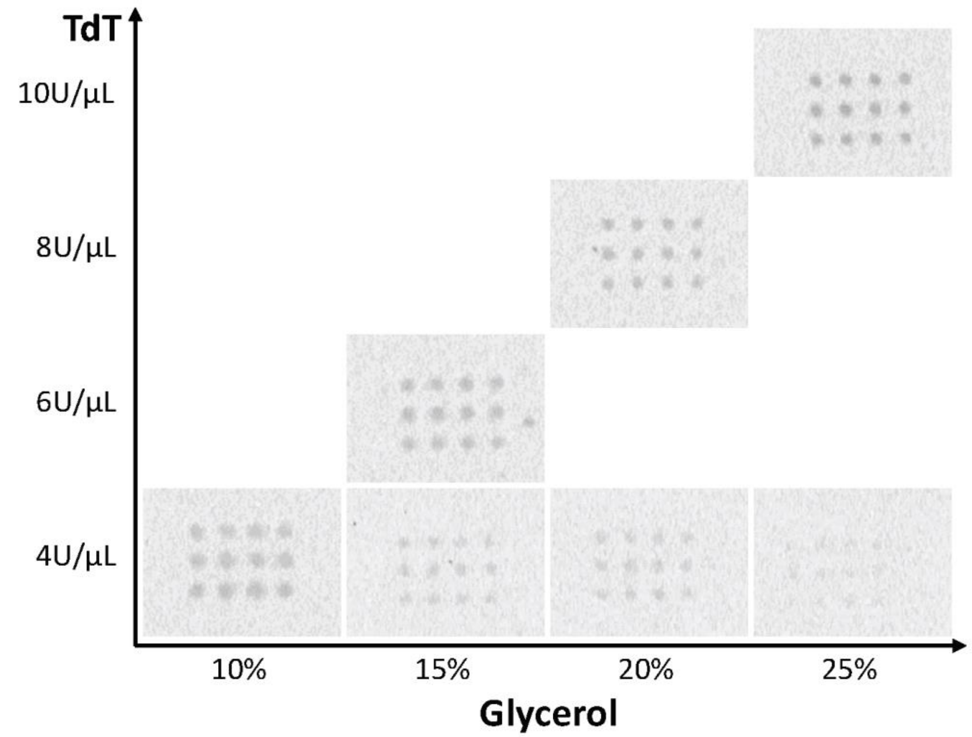


**Supplementary Figure 3:** A comparison of glycerol percentage and concentration of TdT in the synthesis master mix. Increasing the glycerol percentage leads to slower enzyme kinetics, but can be overcome by increasing the total concentration of TdT. The concentration of TdT is defined in units per volume. One unit of TdT catalyzes the incorporation of 1 nmol of deoxythymidylate into a polynucleotide fraction in 60 min at 37 °C (Thermo). However, all reactions were performed at room temperature.

**S4a** **S4b**

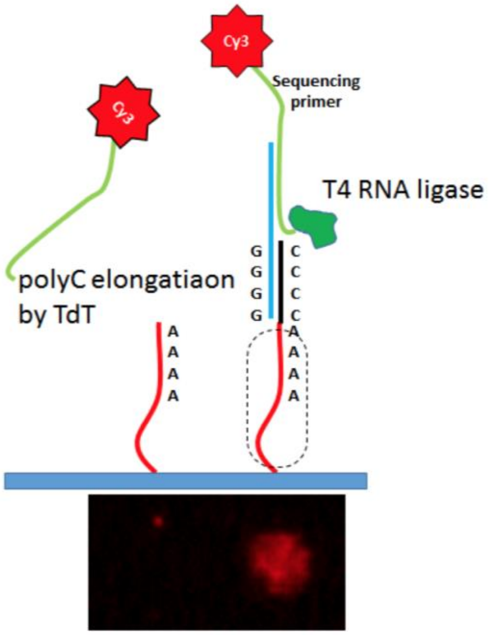


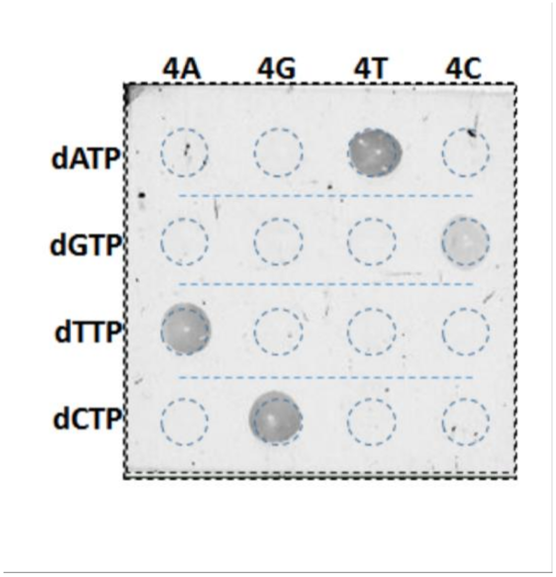


**Supplementary Figure 4: a,** Specific splint-end ligation mechanism for visualization of oligonucleotide synthesis. Surface bound oligonucleotides first undergo a final 3’- extension using the “C” nucleotide by TdT. A ligation master mix containing T4 RNA Ligase, a splint oligonucleotide, and a 3’-Cy3 labeled probe is then incubated with the surface-bound oligonucleotide to attach the probe. After thorough washing, surface bound oligonucleotides can be visualized with fluorescence imaging. An extension of a minimum 4 nucleotides is required for probe attachment using the splint oligonucleotide. Ligation of PCR or NGS adaptors can be attached to surface-bound oligonucleotide in the same manner. **b,** Final extension can be performed with any of the natural nucleotide bases and is not restricted to “C”. For example, a final extension with dATP, requires a splint with a set of “T”. This is demonstrated for each nucleotide extension and splint oligonucleotide combination. Dark spots indicate successful ligation of Cy3 probe.

| Index | Template |
|---|---|
| 0 | GAGACACT |
| 1 | GAGTATAG |
| 2 | GAGCATGA |
| 3 | GACTATAG |
| 4 | GACGCATC |
| 5 | GACACGAG |
| 6 | GATCGCGA |
| 7 | GATATGCT |
| 8 | GATGCTAG |
| 9 | GTCTGCAT |
| 10 | GTCGATAG |
| 11 | GTCATGCA |

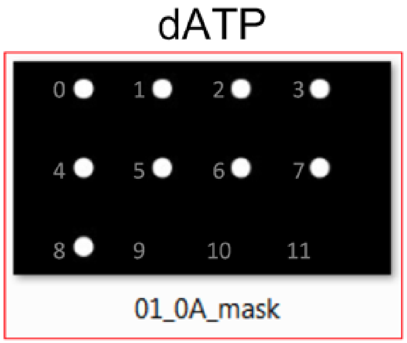


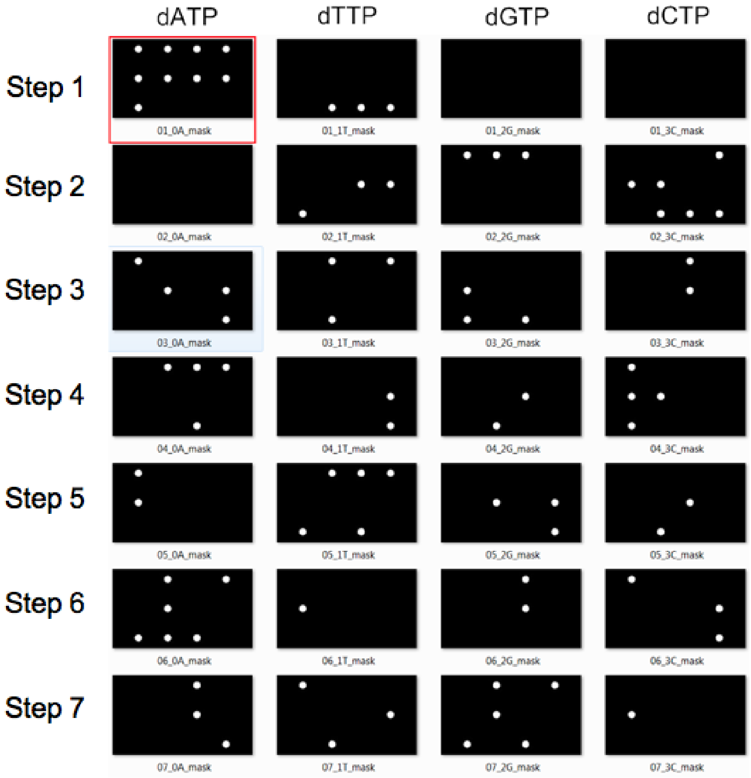


**Supplementary Figure 5:** An overview of all 29 dynamic masks needed to optimally synthesize the 12 DNA oligonucleotide sequences encoding the simplified “Overworld Theme” melody using a (3 × 4) pattern with 100 µm ciricular spots in multiplex. Each row of masks represents a single synthesis step, which encompasses the delivery of each natural nucleotide and the illumination pattern necessary for their spatially specific incorporation as individual cycles. For example, in the synthesis step 1, the “A” nucleotide is needed for sequences being synthesized at indices 0 through 8. This is followed by the “T” nucleotide, which is needed at indices 9 through 11. The “G” and “C” nucleotides are not needed so no masks are generated and the DMD does not illuminate any of the spots during synthesis step 1. This process can be altered to accommodate the synthesis of more sequences, nucleotide bases, and the total sequence template length. On Step 8, all spots on the array are illuminated using one mask to perform the final “C” extension (not shown).

## S6a

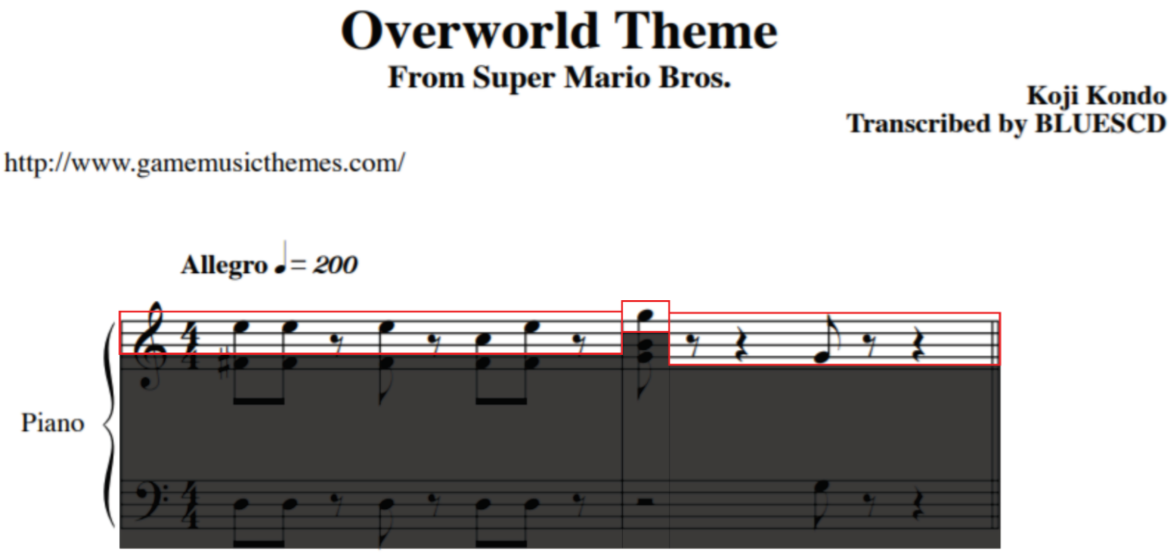


## S6b

| Index | 0 | 1 | 2 | 3 | 4 | 5 | 6 | 7 | 8 | 9 | 10 | 11 |
|---|---|---|---|---|---|---|---|---|---|---|---|---|
| Note | E5 | E5 | $G\#_0$ | E5 | $G\#_0$ | C6 | E6 | $G\#_0$ | G6 | $G\#_0$ | G5 | $G\#_0$ |
| Tempo | ♪ | ♪ | 𝄾 | ♪ | 𝄾 | ♪ | ♪ | 𝄾 | ♪ | 𝄾𝄽 | ♪ | 𝄾𝄽 |

**Supplementary Figure 6: a,** Piano sheet music showing the first two measures of the "Overworld Theme" composed by Koji Kondo for the 1985 Nintendo Entertainment System (NES) video game Super Mario Brothers ™. **b,** Musical notes and their tempos were extracted and indexed from the sheet music to produce a simplified melody to be encoded into DNA sequences as indicated in the table. Rests, where no musical note is played in the melody, were assigned to the note $G\#_0$, which plays at a frequency that is inaudible to the ordinary adult human [2]. The simplified melody is represented on the sheet music with the red box. Grayed-out musical information was not encoded in DNA.

*Sheet music online source:*
*http://www.gamemusicthemes.com/sheetmusic/nintendo/supermariobros/overworldtheme/index.html*

## S7a

#### Original Midi Chart

| Octave | Note Numbers | | | | | | | | | | | |
|---|---|---|---|---|---|---|---|---|---|---|---|---|
| | C | C# | D | D# | E | F | F# | G | G# | A | A# | B |
| -1 | 0 | 1 | 2 | 3 | 4 | 5 | 6 | 7 | 8 | 9 | 10 | 11 |
| 0 | 12 | 13 | 14 | 15 | 16 | 17 | 18 | 19 | 20 | 21 | 22 | 23 |
| 1 | 24 | 25 | 26 | 27 | 28 | 29 | 30 | 31 | 32 | 33 | 34 | 35 |
| 2 | 36 | 37 | 38 | 39 | 40 | 41 | 42 | 43 | 44 | 45 | 46 | 47 |
| 3 | 48 | 49 | 50 | 51 | 52 | 53 | 54 | 55 | 56 | 57 | 58 | 59 |
| 4 | **60** | **61** | **62** | **63** | **64** | **65** | **66** | **67** | **68** | **69** | **70** | **71** |
| 5 | **72** | **73** | **74** | **75** | **76** | **77** | **78** | **79** | **80** | **81** | **82** | **83** |
| 6 | **84** | **85** | 86 | 87 | 88 | 89 | 90 | 91 | 92 | 93 | 94 | 95 |
| 7 | 96 | 97 | 98 | 99 | 100 | 101 | 102 | 103 | 104 | 105 | 106 | 107 |
| 8 | 108 | 109 | 110 | 111 | 112 | 113 | 114 | 115 | 116 | 117 | 118 | 119 |
| 9 | 120 | 121 | 122 | 123 | 124 | 125 | 126 | 127 | | | | |

**Subtract default note number (60)**

#### Modified Midi Chart

| Octave | Note Numbers | | | | | | | | | | | |
|---|---|---|---|---|---|---|---|---|---|---|---|---|
| | C | C# | D | D# | E | F | F# | G | G# | A | A# | B |
| -1 | -60 | -59 | -58 | -57 | -56 | -55 | -54 | -53 | -52 | -51 | -50 | -49 |
| 0 | -48 | -47 | -46 | -45 | -44 | -43 | -42 | -41 | -40 | -39 | -38 | -37 |
| 1 | -36 | -35 | -34 | -33 | -32 | -31 | -30 | -29 | -28 | -27 | -26 | -25 |
| 2 | -24 | -23 | -22 | -21 | -20 | -19 | -18 | -17 | -16 | -15 | -14 | -13 |
| 3 | -12 | -11 | -10 | -9 | -8 | -7 | -6 | -5 | -4 | -3 | -2 | -1 |
| 4 | **0** | **1** | **2** | **3** | **4** | **5** | **6** | **7** | **8** | **9** | **10** | **11** |
| 5 | **12** | **13** | **14** | **15** | **16** | **17** | **18** | **19** | **20** | **21** | **22** | **23** |
| 6 | **24** | **25** | 26 | 27 | 28 | 29 | 30 | 31 | 32 | 33 | 34 | 35 |
| 7 | 36 | 37 | 38 | 39 | 40 | 41 | 42 | 43 | 44 | 45 | 46 | 47 |
| 8 | 48 | 49 | 50 | 51 | 52 | 53 | 54 | 55 | 56 | 57 | 58 | 59 |
| 9 | 60 | 61 | 62 | 63 | 64 | 65 | 66 | 67 | -60 | -60 | -60 | -60 |

## S7b

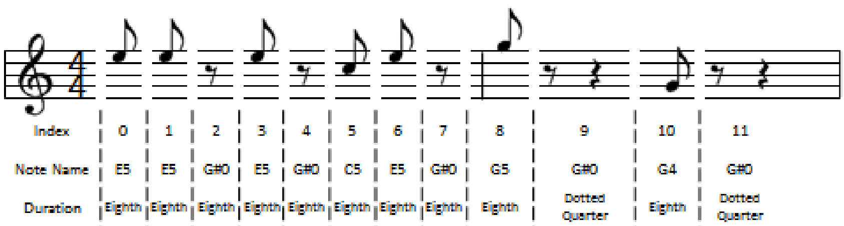


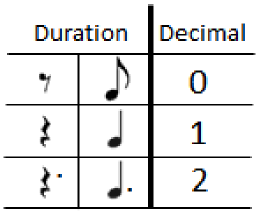

## S7c

| Index (0/11) Ternary form | Note name | Note number (# - default octave 60) | Note number (Ternary form) | Duration | Ternary data | Template Sequence (G start) |
|---|---|---|---|---|---|---|
| 000 | E5 | 76 (16) | 121 | 0 (1X eighth note) | 0001210 | GAGACACT |

**Supplementary Figure 7: a,** The indexed notes from the simplified melody are assigned a note number based on a modified Musical Instrument Digital Information (MIDI) note chart, which indicates both the note and the octave at which it is played at. **b,** These numbers are converted into ternary and combined with the ternary forms of the index and duration number assignments for the note, yielding a 7 digit sequence of numbers. **c,** For example, the first note in the simplified melody is E5 with a quarter note duration. This yields the ternary data **000**1210, where **000** indicates that it is the first note to be played, **121** indicates that the note is E at the 5th octave, and **0** indicates that it should be played for the duration of one eighth note. From this, a template DNA sequence is mapped using previously described methods [1].

*Source for MIDI conversion in MatLab* $^{TM}$*:*
*https://www.mathworks.com/help/audio/ref/midimsg.html*

| | | | | | Ternary | | | | |
|---|---|---|---|---|---|---|---|---|---|
| Index | Note name | Note number | Note # -default octave | Duration | Index | Note number | Duration | Data | Template |
| 0 | E5 | 76 | 16 | Eighth | 000 | 121 | 0 | 0001210 | GAGACACT |
| 1 | E5 | 76 | 16 | Eighth | 001 | 121 | 0 | 0011210 | GAGTATAG |
| 2 | G#0 | 20 (86) | 26 | Eighth | 002 | 222 | 0 | 0022220 | GAGCATGA |
| 3 | E5 | 76 | 16 | Eighth | 010 | 121 | 0 | 0101210 | GACTATAG |
| 4 | G#0 | 20 (86) | 26 | Eighth | 011 | 222 | 0 | 0112220 | GACGCATC |
| 5 | C5 | 72 | 12 | Eighth | 012 | 110 | 0 | 0121100 | GACACGAG |
| 6 | E5 | 76 | 16 | Eighth | 020 | 121 | 0 | 0201210 | GATCGCGA |
| 7 | G#0 | 20 (86) | 26 | Eighth | 021 | 222 | 0 | 0212220 | GATATGCT |
| 8 | G5 | 79 | 19 | Eighth | 022 | 201 | 0 | 0222010 | GATGCTAG |
| 9 | G#0 | 20 (86) | 26 | Eighth+Quarter | 100 | 222 | 2 | 1002222 | GTCTGCAT |
| 10 | G4 | 67 | 7 | Eighth | 101 | 021 | 0 | 1010210 | GTCGATAG |
| 11 | G#0 | 20 (86) | 26 | Eighth+Quarter | 102 | 222 | 2 | 1022222 | GTCATGCA |

**Supplementary Figure 8:** A table outlining the fully converted "Overworld Theme" simplified musical melody to ternary information and then mapped to a DNA template sequence to be synthesized in multiplex. Note that all mapped DNA template sequences start with the "G" nucleobase, this is the 3'- end of the surface-bound initiator oligonucleotide that all sequences will be synthesized from.

**S9a**

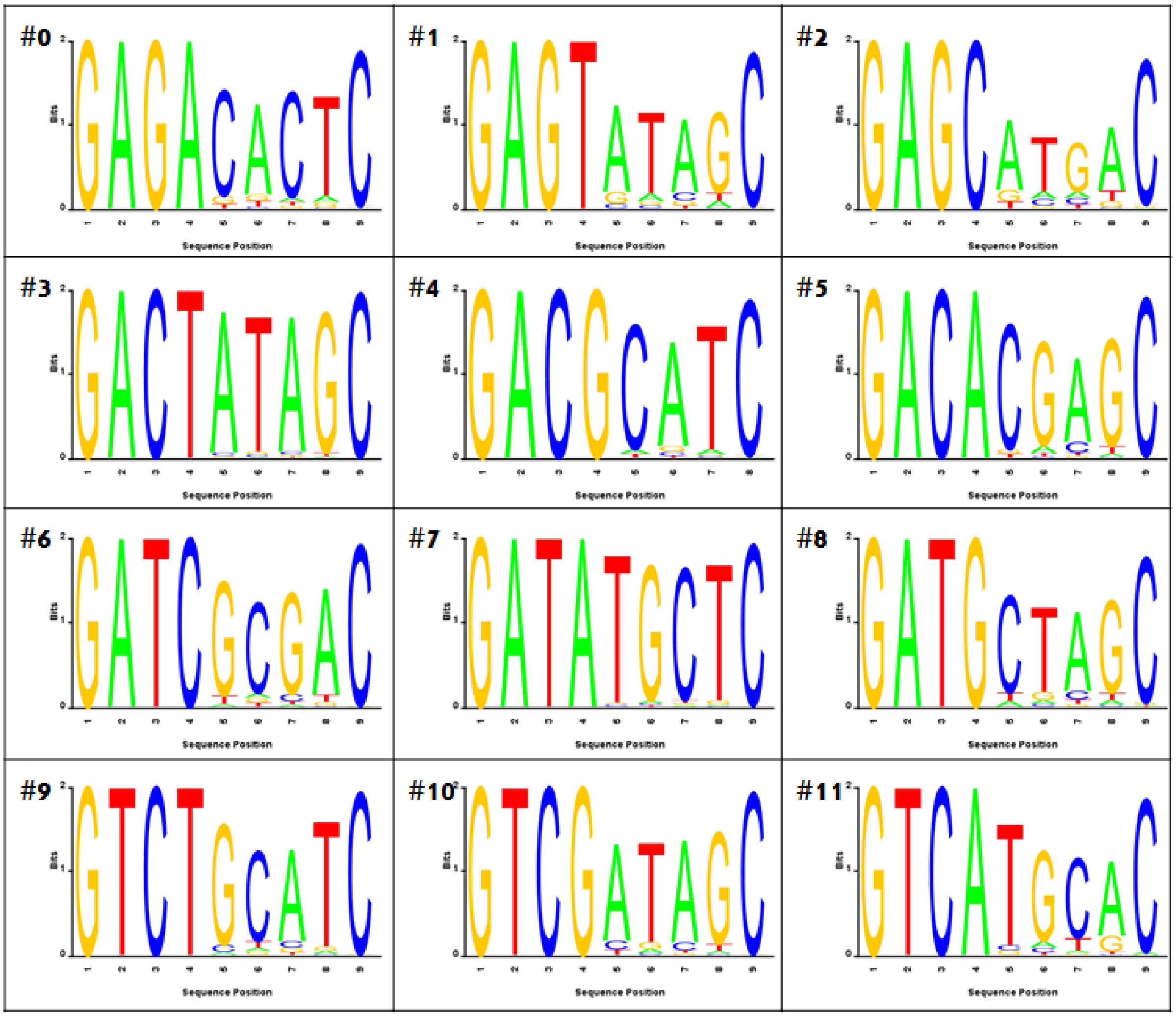

# S9b

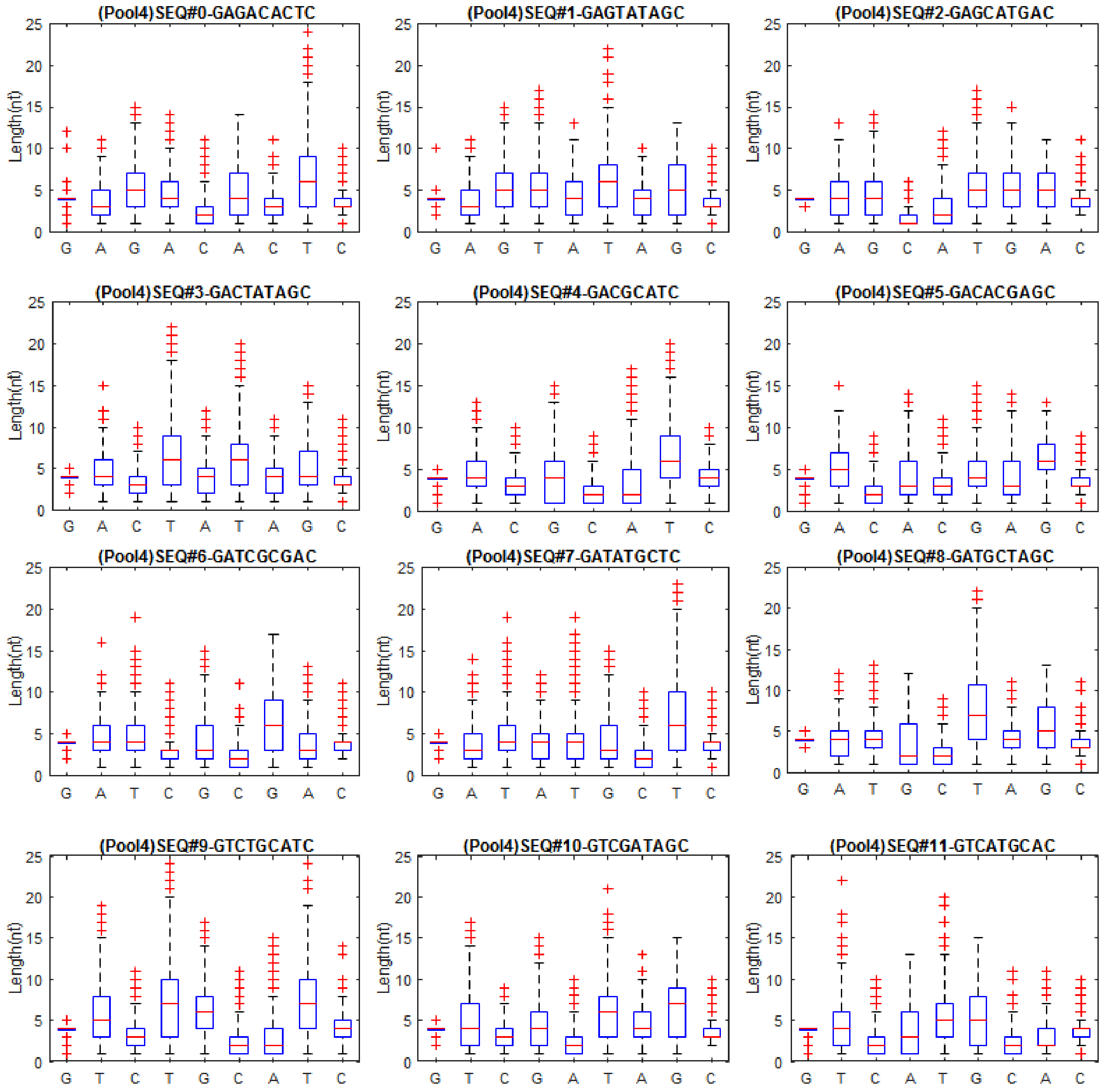
(Pool4)SEQ#0-GAGACACTC
(Pool4)SEQ#1-GAGTATAGC
(Pool4)SEQ#2-GAGCATGAC
(Pool4)SEQ#3-GACTATAGC
(Pool4)SEQ#4-GACGCATC
(Pool4)SEQ#5-GACACGAGC
(Pool4)SEQ#6-GATCGCGAC
(Pool4)SEQ#7-GATATGCTC
(Pool4)SEQ#8-GATGCTAGC
(Pool4)SEQ#9-GTCTGCATC
(Pool4)SEQ#10-GTCGATAGC
(Pool4)SEQ#11-GTCATGCAC
Length(nt)

## S9c

| | |
|---|---|
| **Total reads** | **216252** |
| **Sequence with adaptor** | **100760** |
| **Perfect transition match** | **14335** |

| Sequence | GAGACACTC | GAGTATAGC | GAGCATGAC | GACTATAGC | GACGCATC | GACACGAGC | GATCGCGAC | GATATGCTC | GATGCTAGC | GTCTGCATC | GTCGATAGC | GTCATGCAC |
|---|---|---|---|---|---|---|---|---|---|---|---|---|
| Sequence index | 0 | 1 | 2 | 3 | 4 | 5 | 6 | 7 | 8 | 9 | 10 | 11 |
| Filter 1,2,3 (# of transition, index) | 1742 | 605 | 426 | 2357 | 2268 | 1053 | 1182 | 2668 | 536 | 2344 | 878 | 682 |
| Filter 4 (perfect match) | 1335 | 466 | 303 | 2182 | 2014 | 868 | 996 | 2492 | 423 | 1993 | 739 | 524 |

**Supplementary Figure 9:** In-depth analysis for Illumina miSeq sequencing. **a,** Sequence logo representation of all 12 strands after *in silico* filter 1, 2, and 3. **b,** Box plot statistical information for the extension length distribution for each base transition for all perfectly matched (*in silico* filter 1, 2, 3, and 4) 12 oligonucleotides synthesized in multiplex. Each SEQ represents an individual subset/index and DNA oligonucleotide. **c,** Raw sequencing data was filtered as per specified elsewhere for perfect matches containing all 8 base transitions for each oligonucleotide index. The breakdown for filtering from the raw sequencing data is included.

**S10a**

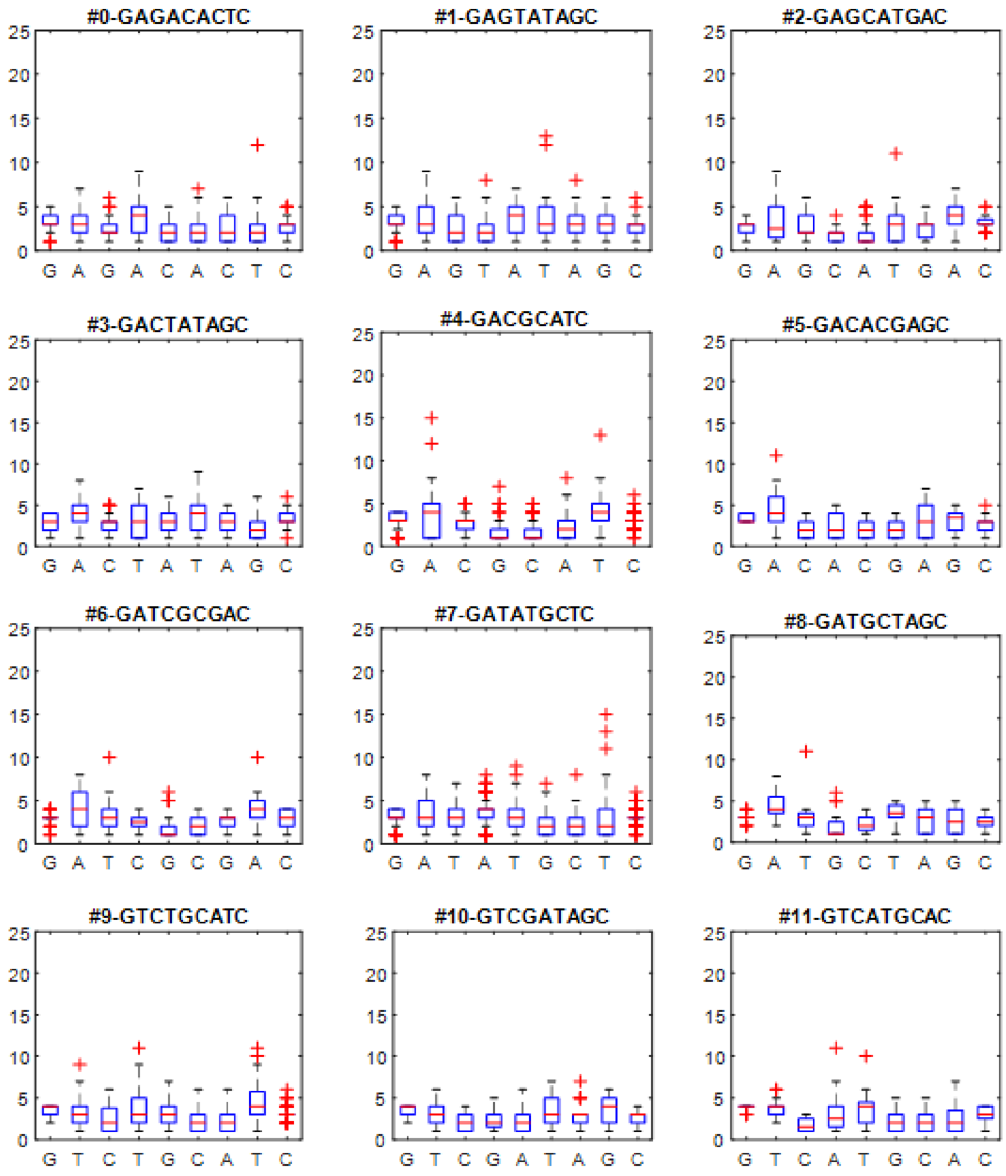

#0-GAGACACTC
#1-GAGTATAGC
#2-GAGCATGAC
#3-GACTATAGC
#4-GACGCATC
#5-GACACGAGC
#6-GATCGCGAC
#7-GATATGCTC
#8-GATGCTAGC
#9-GTCTGCATC
#10-GTCGATAGC
#11-GTCATGCAC

## S10b

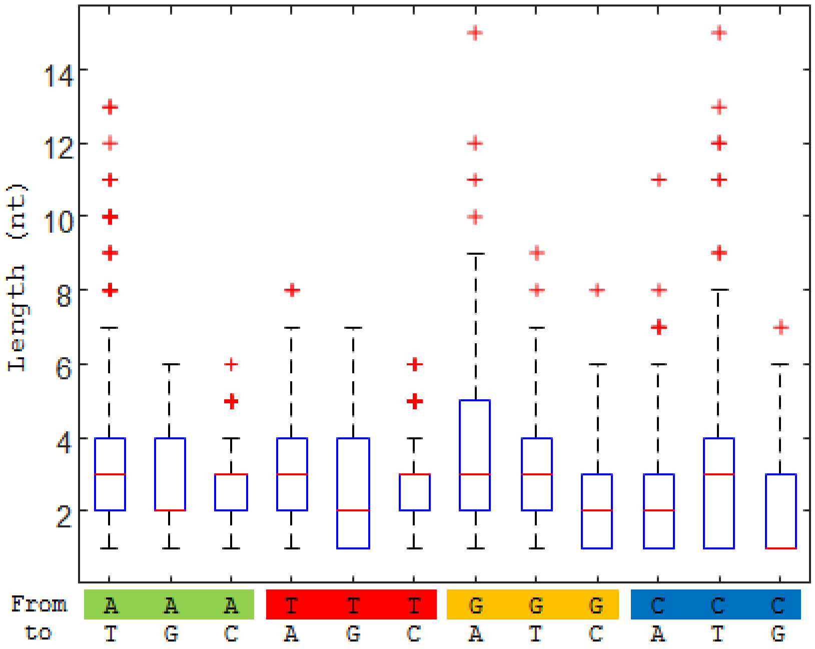


## S10c

| | |
|---|---|
| **Total reads** | **946872** |
| **Sequence with adaptor** | **20014** |
| **Perfect transition match** | **569** |

| Sequence | GAGACACTC | GAGTATAGC | GAGCATGAC | GACTATAGC | GACGCATC | GACACGAGC | GATCGCGAC | GATATGCTC | GATGCTAGC | GTCTGCATC | GTCGATAGC | GTCATGCAC |
|---|---|---|---|---|---|---|---|---|---|---|---|---|
| Sequence index | 0 | 1 | 2 | 3 | 4 | 5 | 6 | 7 | 8 | 9 | 10 | 11 |
| Filter 1,2,3 (# of transition, index) | 252 | 166 | 89 | 86 | 106 | 66 | 74 | 175 | 51 | 112 | 47 | 57 |
| Filter 4 (perfect match) | 46 | 78 | 32 | 56 | 77 | 18 | 22 | 121 | 16 | 63 | 24 | 16 |

**Supplementary Figure 10:** In-depth analysis using Oxford nanopore sequencing. **a,** Box plot statistical information for the extension length distribution for each base transition for all perfectly matched (*in silico* filter 1, 2, 3, and 4) 12 oligonucleotides synthesized in multiplex. Each SEQ represents an individual subset/index and DNA oligonucleotide. **b,** Statistics for the extension length distribution for all possible transitions from the entire array. **c,** Raw sequencing data was filtered as per specified elsewhere for perfect matches containing all 8 base transitions for each oligonucleotide index. The breakdown for filtering from the raw sequencing data is included.

### References

1. Lee, H. H., Kalhor, R., Goela, N., Bolot, J. & Church, G. M. Terminator-free template-independent enzymatic DNA synthesis for digital information storage. *Nat. Commun.* **10**, 2383 (2019).
2. Rosen, S. & Howell, P. *Signals and Systems for Speech and Hearing*. (BRILL, 2011).
